# Impact of neurite alignment on organelle motion

**DOI:** 10.1101/2021.07.26.453779

**Authors:** Maria Mytiliniou, Joeri A. J. Wondergem, Thomas Schmidt, Doris Heinrich

**Affiliations:** Leiden Institute of Physics, Huygens-Kamerlingh Onnes Laboratory, Leiden University, Leiden, the Netherlands; Institute for Bioprocessing and Analytical Measurement Techniques, Rosenhof, 37308 Heilbad Heiligenstadt, Germany; Faculty for Mathematics and Natural Sciences, Technische Universität Ilmenau, Ilmenau, Germany; Fraunhofer Institute for Silicate Research ISC, Würzburg, Germany

## Abstract

Intracellular transport is pivotal for cell growth and survival. Malfunctions in this process have been associated with devastating neurodegenerative diseases, posing a need for deeper understanding of the involved mechanisms. Here, we used an experimental methodology that lead neurites of differentiated PC12 cells in either of two configurations: an one-dimensional, where the neurites align along lines, or a two-dimensional configuration, where the neurites adopt a random orientation and shape on a flat substrate. We subsequently monitored the motion of functional organelles, the lysosomes, inside the neurites. Implementing a time-resolved analysis of the mean-squared displacement, we quantitatively characterized distinct motion modes of the lysosomes. Our results indicate that neurite alignment gives rise to faster diffusive and super-diffusive lysosomal motion in comparison to the situation where the neurites are randomly oriented. After inducing lysosome swelling through an osmotic challenge by sucrose, we confirmed the predicted slowdown in diffusive mobility. Surprisingly we found that the swelling-induced mobility change affected each of the (sub-/super-) diffusive motion modes differently and depended on the alignment configuration of the neurites. Our findings imply that intracellular transport is significantly and robustly dependent on cell morphology, which might be in part controlled by the extracellular matrix.

## Introduction

One critical function not only for the growth and maintenance of homeostasis, but also for the survival of a cell, is the transport of proteins, molecules, organelles and debris to specific locations within the cell. This allocation is of tremendous significance, especially for neuronal cells due to the extreme size of their axons; for instance, axons of human motor neurons can reach the length of one meter, starting from the brain and extending till the end of the spine. Defects in the process of intracellular transport have long been associated with human diseases (Aridor and Lisa A. Hannan 2000; Aridor and Lisa A Hannan 2002; Sleigh et al. 2019; Appert-Rolland, Ebbinghaus, and Santen 2013; De Vos and Hafezparast 2017), but whether such defects are the cause or the consequence of pathological phenotypes, is still under debate in several cases (Moloney, Winter, and Verhaagen 2014; Prior et al. 2017).

Intracellular distribution of molecules and organelles is achieved mainly via two mechanisms: passive diffusion and active, motor-driven, transport along microtubules and actin filaments (Vale 2003). Cytoskeletal components, organelles and molecules crowd the cytoplasm, thereby hindering or enhancing passive diffusion, thus leading to sub-diffusive and super-diffusive intracellular motion (Götz et al. 2015; Otten et al. 2012; Witzel et al. 2019; S Mogre, Brown, and Koslover 2020). In order to characterize intracellular dynamics and extract values such as the velocity of the motor-mediated transport or the diffusion coefficient of passive motion, several models have been implemented (Bressloff and Newby 2013; Briane, Kervrann, and Vimond 2018; Norregaard et al. 2017).

A frequently used approach exploits the Mean Squared Displacement (MSD) plotted as a function of lag time and subsequently fitted with a power law in the form of ~2dD*τ*^*α*^, where d is the dimensionality, D the diffusion coefficient and *τ* the lag time (Gal, Lechtman-Goldstein, and Weihs 2013; Grady et al. 2017). The characteristic exponent (*α*) value reveals the type of motion, differentiating among Brownian diffusion (*α* = 1), from now on referred to as diffusion, super-diffusion (*α* > 1) and sub-diffusion (*α* < 1). In the case of complex motion, such as intracellular transport, comprising a combination of alternating phases of sub-diffusion, free diffusion and motor-driven active transport, motion discriminating algorithms are essential to avoid averaging out dynamic information. Thus, local-MSD (lMSD) analysis can be performed instead of fitting the entire MSD curve, providing time-resolved information of the motion states within a trajectory. This analysis implements a rolling window over the entire trajectory, thereby characterizing each data point with the *α* exponent value (Arcizet et al. 2008; Mahowald, Arcizet, and Heinrich 2009; Otten et al. 2012; Dupont et al. 2013; Götz et al. 2015).

Although the MSD analysis is a well-established tool, a reliable and less noisy MSD curve requires many data points and long trajectories which, in the case of biological data, can be challenging to acquire (Michalet 2010). Additionally, selection of the model to fit is not always straightforward, especially in the instance of complex data with multiple underlying processes (Türkcan and Masson 2013). Furthermore, the MSD curve is sensitive to experimental parameters such as the acquisition frame rate and the size of the imaged particle (Gal, Lechtman-Goldstein, and Weihs 2013).

Hence, the use of an additional method, the van Hove distribution (Van Hove 1954), for the analysis of intracellular data can counterweight the drawbacks described above and provide a more in-depth view of intracellular transport than that gained from the MSD analysis alone. The van Hove distribution, alternatively called Jump Distance Distribution (JDD) was initially used for particle scattering experiments (De Bar 1963; Dahlborg, Gudowski, and Davidovic 1989). The JDD is plotted for a specific lag time and depicts the Euclidian displacement distribution of the observed particles within the given time lag, in the form of a Probability Distribution Function (PDF). Thus, this distribution reveals the displacement-dependent structure which is, otherwise, “hidden” in a single, averaged data point, for the respective lag time, of the MSD curve.

Already in 1997, Schütz et al. showed that by fitting the probability distribution of the squared displacements of single-molecules moving on membranes, one can extract individual diffusion constants and fractions of multiple-component samples (Schütz, Schindler, and Schmidt 1997). Along the same lines, Kues et al. analyzed single-molecule motion inside cell nuclei, distinguishing among three mobility states (Kues, Peters, and Kubitscheck 2001). Over time, application of this analysis for Brownian motion in fluids (Hopkins et al. 2010) and in - actual or simulated - biological systems increased (Ghosh et al. 2016; Menssen and Mani 2019; Grady et al. 2017; Bhowmik, Tah, and Karmakar 2018; Witzel et al. 2019), establishing it as a powerful tool for characterization of complex biological trajectories.

Here, we set out to characterize trajectories of lysosomes inside neurites of differentiated PC12 cells (Lloyd A Greene and Tischler 1976; Lloyd A Greene 1978; Lloyd A. Greene et al. 1987), commonly used as neuronal model (Wang et al. 2015; Christen et al. 2017; Pelzl et al. 2009). Neurites are the precursors of dendrites and axons in immature neurons (Leterrier, Dubey, and Roy 2017) hence, motion analysis within neurites can provide significant insight into axonal transport. Lysosomes are organelles that play a vital role in the autophagy pathway of cells and exhibit both diffusive motion and active transport via dynein and kinesin motor proteins along microtubules. Not only the autophagy pathway in general (Nixon 2013; Menzies, Fleming, and Rubinsztein 2015; Ramesh and Pandey 2017), but also the motion of lysosomes specifically appear to be implicated with neurodegenerative diseases and cancer (Lawrence and Zoncu 2019; Amick and Ferguson 2017; Burk and Pasterkamp 2019; Oyarzún et al. 2019).

We show that neurite alignment, achieved via chemical surface patterning, results in faster diffusive and super-diffusive lysosomal motion in comparison to the case where the neurites adopt a random orientation. Moreover, we introduce a perturbation in the cellular environment via incubation with sucrose and confirm experimentally that the sucrose induces lysosomal enlargement, which leads to a proportionate decrease in the diffusion coefficient. Implementing lMSD analysis, we identify and extract the trajectory parts that belong into each of three classes of motion, namely sub-diffusive, diffusive and super-diffusive. By collectively analyzing the data points of the respective class, we gain quantitative insights for each motion mode. Our findings indicate that the incubation with sucrose results in a different effect on each motion mode of the organelles, and this also depends on the configuration of the neurites within which the motion occurs.

## Results

We set out to characterize intracellular organelle transport and to compare motion features under two distinct neurite configurations. We investigated the motion of lysosomes inside neurites of differentiated PC12 cells, when those adopt a random orientation on the two-dimensional culture surface, versus when they are prompted to adhere to an one-dimensional configuration by means of chemical surface patterning.

### microscale Plasma - Initiated Patterning (*μ*PIP) of Laminin guides the neurites of differentiated PC12 cells along 2 *μ*m-wide lines

We achieved one-dimensional neurite alignment by selective protein deposition on the cell substrate. The steps followed during the *μ*PIP are schematically shown in Fig. 1.C-F. A polydimethylsiloxane (PDMS) mask bearing a ladder-shape pattern was used. Scanning-electron microscopy (SEM) images of the PDMS mask are shown in Fig. 1.A and B. The mask was inverted and pressed onto the cell substrate. The assay was then exposed to air plasma (Fig. 1.C), thereby altering the surface charge of the areas exposed via the mask and thus increasing their hydrophilicity. The rest of the surface, covered by the adhered PDMS mask, remained in its original hydrophobic state. Subsequent incubation with Pluronic F127 and Laminin (Fig. 1.D and E, respectively), resulted in 2*μ*m-wide Laminin lines, alternating with 18*μ*m-wide Pluronic F127-coated stripes (Fig. 1.F).

**Figure 1:**
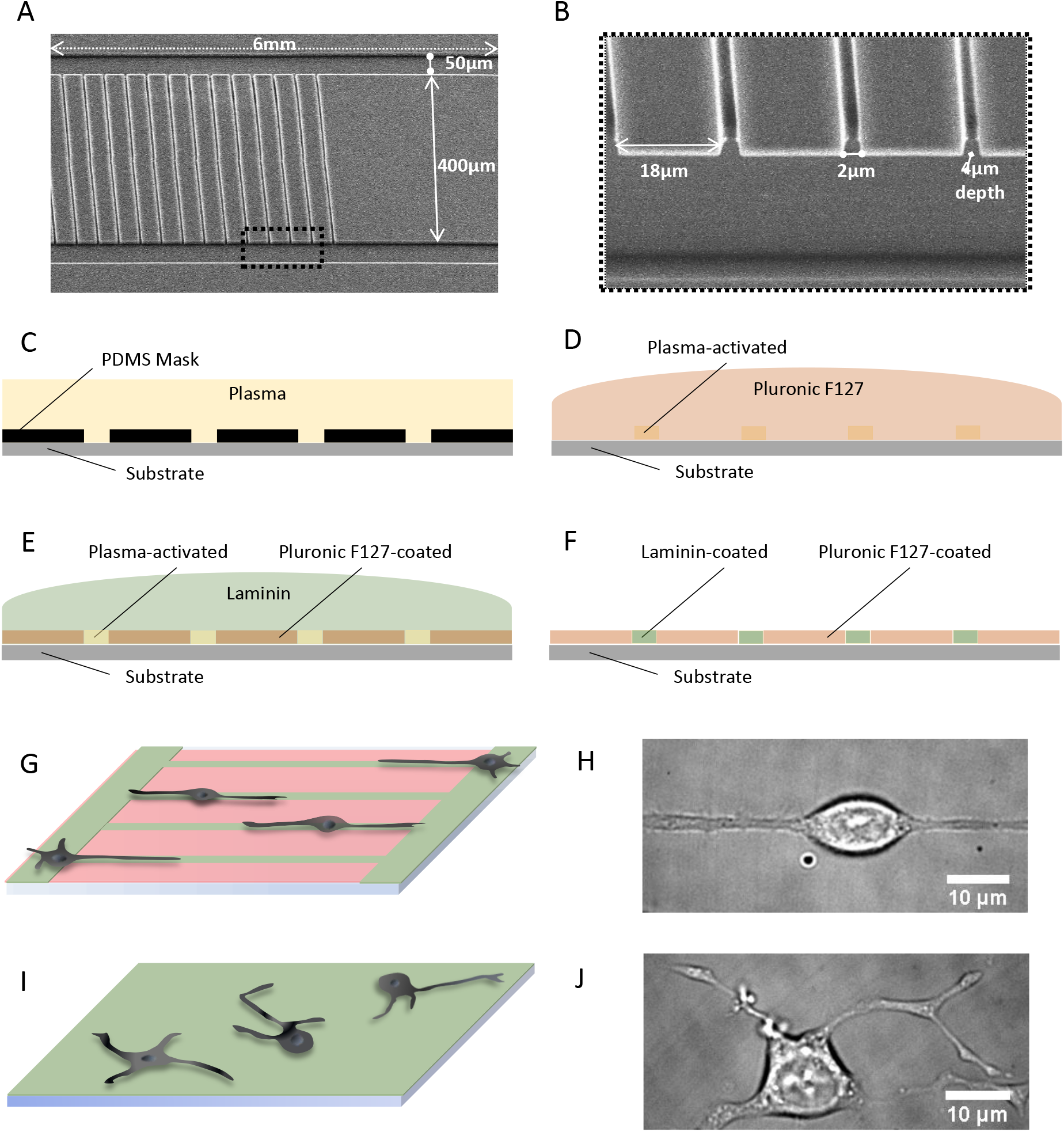
Laminin *μ*PIP for neurite guidance. (A) Scanning electron microscopy (SEM) image of the PDMS mask used for the plasma-initiated patterning. The structure consisted of two side lines, 50*μ*m-wide and 6mm long, with 400*μ*m distance in-between. The central 2mm of the 6mm blocks were intersected by 2*μ*m-wide lines, repeated every 18*μ*m. The depth of the mask was 4*μ*m. (B) Close-up of the area indicated by the black dotted-line square in (A). (C-F) Steps followed for the *μ*PIP: (C) The PDMS mask was placed on the substrate, and the assay was exposed to air plasma. (D) The PDMS mask was removed and the substrate was submerged in Pluronic F127 which adsorbed to the plasma-protected areas. (E) The dish was immersed in Laminin, which adhered to the plasma-activated areas. (F) The resulting pattern of Laminin-coated lines surrounded by Pluronic-covered regions. (G) Schematic of the patterned substrate. Green color represents the ECM protein (Laminin) and red color the Pluronic F127. (H) Representative bright-field image of a differentiated PC12 cell on the patterned substrate, with its neurites aligned along the line. (I) Schematic of the un-patterned substrate, coated with Laminin (green color). (J) Representative bright-field image of a differentiated PC12 cell with the neurites randomly oriented on two-dimensions.

In order to obtain the second neurite geometry, the entire substrate was coated with the extracellular matrix (ECM) protein. PC12 cells were allowed to adhere on both substrates, as schematically shown in Fig. 1.G and I. Subsequently, the cells were differentiated to stimulate neurite growth (see Materials and Methods). Depending on the substrate used, patterned or un-patterned, the neurites either aligned along the lines or grew randomly on the 2D surface. Representative cells of both configurations are shown in Fig. 1.H and J, respectively.

### Sucrose induces lysosomal enlargement in differentiated PC12 cells

Next, we wondered whether a perturbation in the cellular environment, that associates with lysosomes, could be deciphered by studying their motion and, if that was the case, whether the effect would differ for the two different neurite configurations. To investigate that question, we employed sucrose-induced swelling of lysosomes. Sucrose has long been known to trigger swelling of lysosomes; it enters the cytoplasm by pinocytosis but can not be degraded by lysosomal enzymes, thus causing osmotic pressure alterations in lysosomes, which in turn, attempting to maintain osmotic balance, allow water influx, thus swelling (Warburton and Wynn 1976). Lysosome enlargement, along with its effect on lysosomal transport has been quantified in BS-C-1 monkey kidney epithelial cells (Bandyopadhyay et al. 2014).

Along the same lines, we incubated the differentiated PC12 cells with sucrose prior to data acquisition. Representative images of fluorescent lysosomes inside differentiated PC12 cells in normal (control) media and in media containing sucrose are displayed in Fig. 2. A small effect on the size of the lysosomes can be observed by visual inspection. To quantify this, we measured the diameters of lysosomes for the two conditions and the distributions of the values, are shown in Fig. 2.E. The mean diameter was found to be equal to:

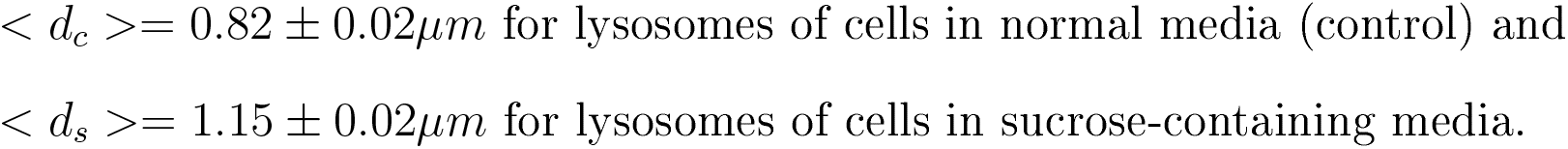

resulting in an increase of 0.33*μ*m for the average lysosome diameter caused by incubation with sucrose.

**Figure 2:**
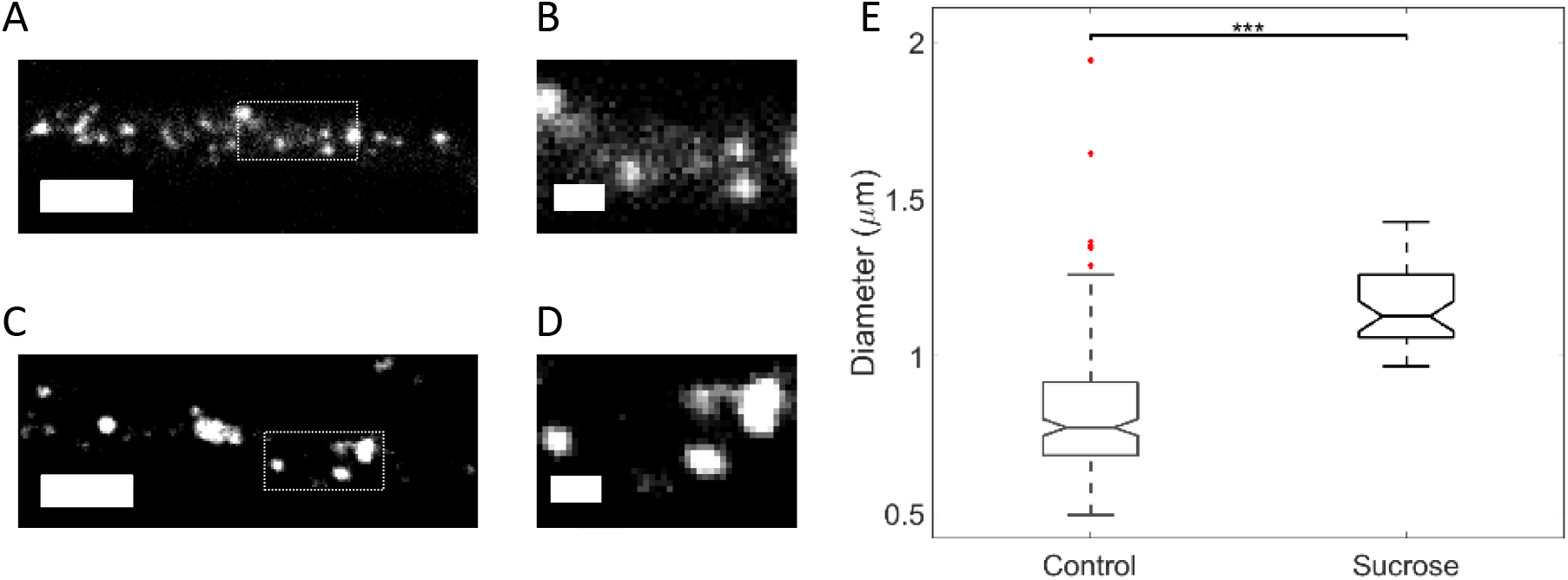
Sucrose induces increase of lysosome diameter. (A, B) Fluorescent lysosomes of a cell in media without sucrose (control). (C, D) Fluorescent lysosomes of a cell in media with sucrose. Scale bar in (A, C) is 5*μ*m and the square indicates the location shown in higher magnification in (B, D), respectively, with scale bar 1*μ*m. (E) Boxplot of lysosomes diameters for differentiated PC12 cells in media without sucrose (control) versus in media with sucrose, with mean values equal to < *d*_*c*_ >= 0.82±0.02*μm* and < *d*_*s*_ >= 1.15±0.02*μm* respectively. The statistical significance between the two means was determined using the Wilcoxon ranksum test; *** corresponds to *p* < 0.001.

### Lysosomes inside aligned neurites exhibit higher displacements

Fluorescently-labeled lysosomes were tracked for up to 30 seconds, inside both neurite configurations. The MSDs and JDD PDFs were calculated for all lysosomal trajectories of each condition (using eq. 5 and 10, respectively), for the x- and y- displacements, and are displayed in Fig. 3. The x-axis coincided with the neurite alignment axis, in the corresponding experimental configuration.

**Figure 3:**
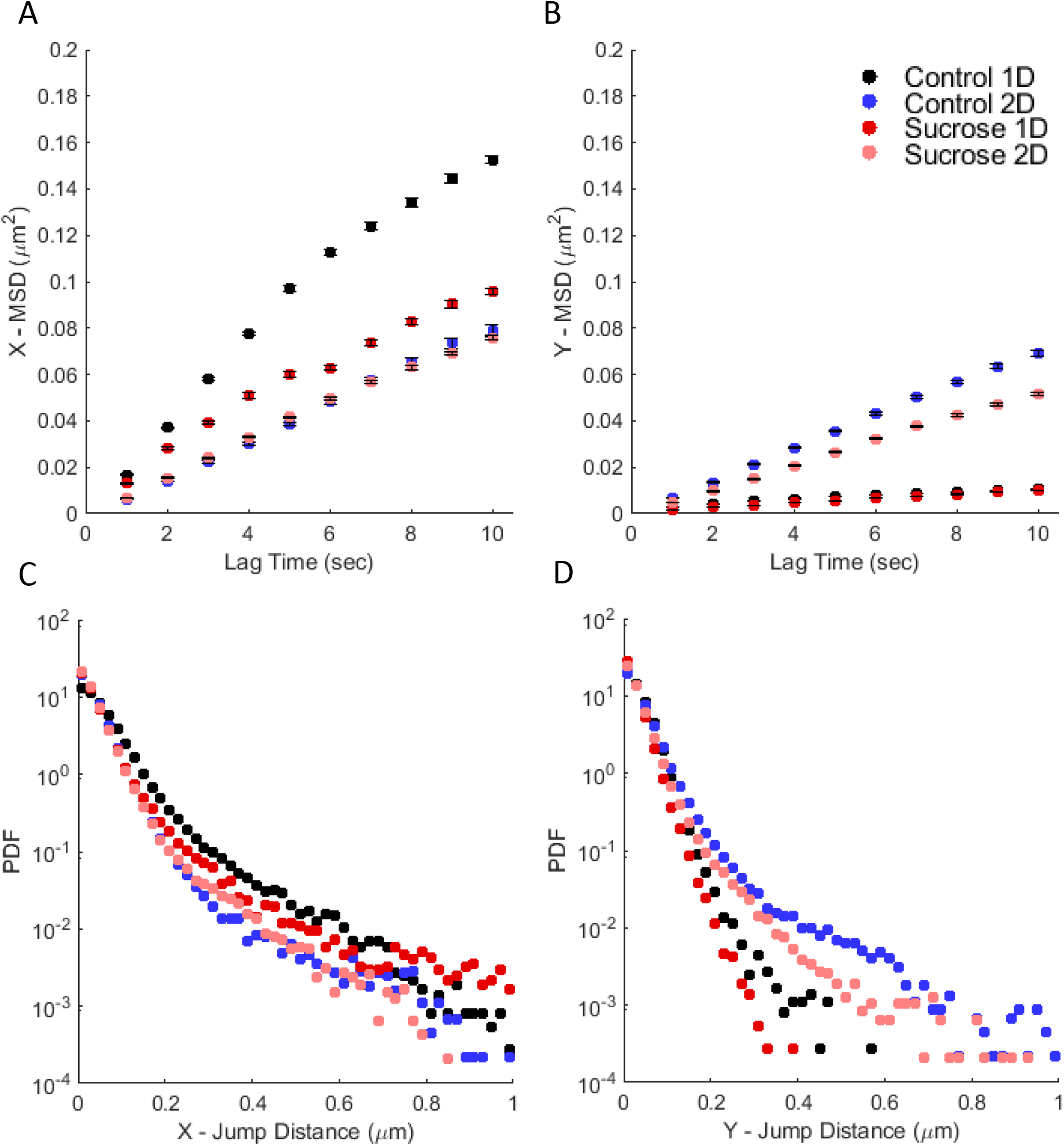
Larger MSD and JDD PDF values observed for lysosomes moving inside aligned neurites. (A) MSD curves along the x- axis and (B) MSD curves along the y- axis of lysosomal trajectories. Data points show time-averaged MSD ± standard error of the mean. (C) JDD PDFs along the x- axis and (D) JDD PDFs along the y- axis of lysosomal trajectories. Color-coding indicates lysosomes inside aligned (1D) or randomly oriented (2D) neurites of differentiated PC12 cells in media without (control) or with sucrose.

As can be observed in Fig. 3.A, the MSD curve along the x- axis exhibits significantly higher values for lysosomes inside aligned neurites, especially for the control condition. It is noteworthy, that even in the presence of the sucrose-induced perturbation, the x-MSD values of lysosomes inside aligned neurites are higher than those of the control condition in randomly oriented neurites. Moreover, the decrease in the displacement observed in the presence of sucrose, is more significant for lysosomes inside aligned neurites, as compared to randomly oriented neurites. These findings are consistent with the JDD PDFs along the x-axis (Fig. 3.C).

The MSD and JDD PDF along the y-axis (Fig. 3.B and D) clearly indicate the underlying neurite alignment of the corresponding experimental condition. On the other hand, the y-MSD and JDD PDF values of lysosomes inside randomly oriented neurites, are similar to those for the x-axis. This result is expected, since there is no directionality preference for lysosomes moving inside randomly oriented neurites.

### Local MSD analysis of lysosomes trajectories distinguishes among sub-diffusive, diffusive and super-diffusive motion modes

The shape of the MSD and JDD PDF curves presented in Fig. 3 indicates that the motion analyzed here consists of more than one type of transport, as explained previously. To characterize each transport mode, we performed a lMSD analysis for every single lysosomal trajectory. Previous studies have implemented this time-resolved analysis, however either distinguishing only between active and passive transport (Ahmed, Williams, et al. 2013; Ahmed and Saif 2014), or without afterwards analyzing collectively the trajectory parts of each motion category (Arcizet et al. 2008; Götz et al. 2015; Mahowald, Arcizet, and Heinrich 2009).

Here, we differentiated among the three modes of lysosomal motion, characterized each trajectory data point, and subsequently analyzed collectively the trajectory parts of each motion type. Fig. 4.A shows the bright-field image of a differentiated cell, with its neurite aligned along the Laminin line, overlayed with recorded trajectories of lysosomes. Fig. 4.B displays the lysosomal trajectory indicated by the dotted black square in Fig. 4.A. The trajectory parts are color-coded, indicating either of sub-diffusive (black), diffusive (blue) and super-diffusive (green) transport. For each mode of motion in this trajectory, the respective MSD curve is displayed in Fig. 4.C. In the same plot, dashed lines indicate theoretical MSD curves of super-diffusion (*α* ~ 1.5) and sub-diffusion (*α* ~ 0.5).

**Figure 4:**
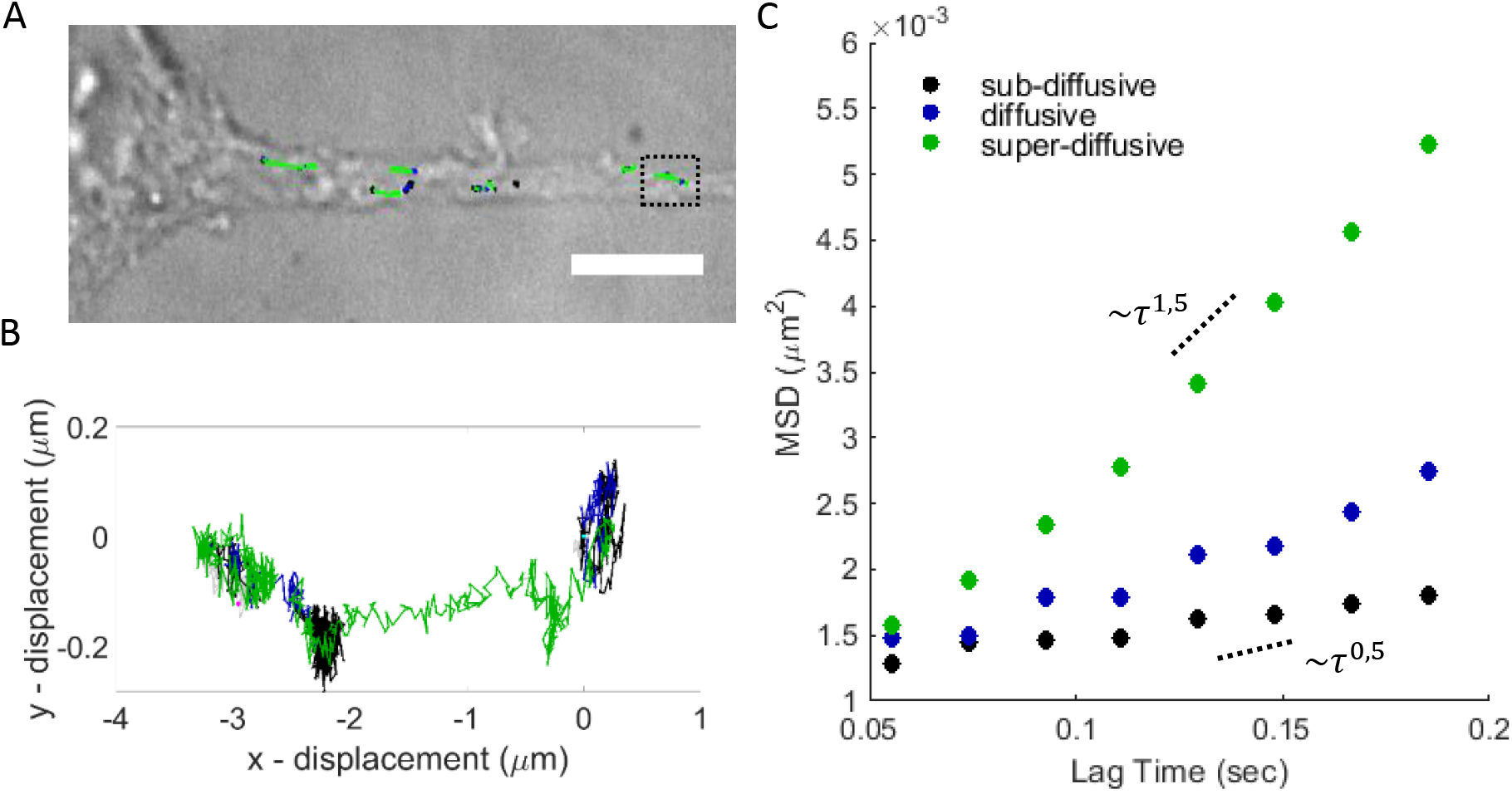
Time-resolved characterization of motion, based on local MSD analysis. (A) Bright-field image of a differentiated PC12 cell with its neurite aligned along the patterned Laminin line. Recorded trajectories of lysosomes are overlayed and color-coded based on the type of motion (sub-diffusive for *α* ≤ 0.9, diffusive for 0.9 ≤ *α* ≤ 1.1 and super-diffusive for *α* ≥ 1.1). The *α* value was determined using the local MSD analysis. Scale bar equals 100*μ*m. (B) Close-up of the trajectory indicated by the black dotted square in (A). (C) Mean-squared displacement of the three motion modes, from the parts comprising the trajectory shown in (B). The dotted lines indicate the theoretical sub-diffusive (*α* ~ 0.5) and super-diffusive (*α* ~ 1.5) MSD curves.

### Neurite alignment associates with more efficient (sub-/ super-) diffusive transport of lysosomes

After characterizing every individual trajectory in a time-resolved manner using the lMSD analysis, we analyzed collectively all trajectory data points of each transport type. The MSD curves of the sub-diffusive, diffusive and super-diffusive parts of lysosomal trajectories in neurites of differentiated PC12 cells for the four experimental conditions, are displayed in Fig. 5. The sub-diffusive MSD curves were fitted using the power law describing anomalous diffusion (eq. 7). To fit the diffusive MSDs, the Brownian motion model was used (eq. 8). Lastly, for the super-diffusive MSDs we implemented the model of Brownian motion with drift (eq. 9) (Briane, Kervrann, and Vimond 2018). The resulting fitting parameters are summarized in Table S.1.

**Figure 5:**
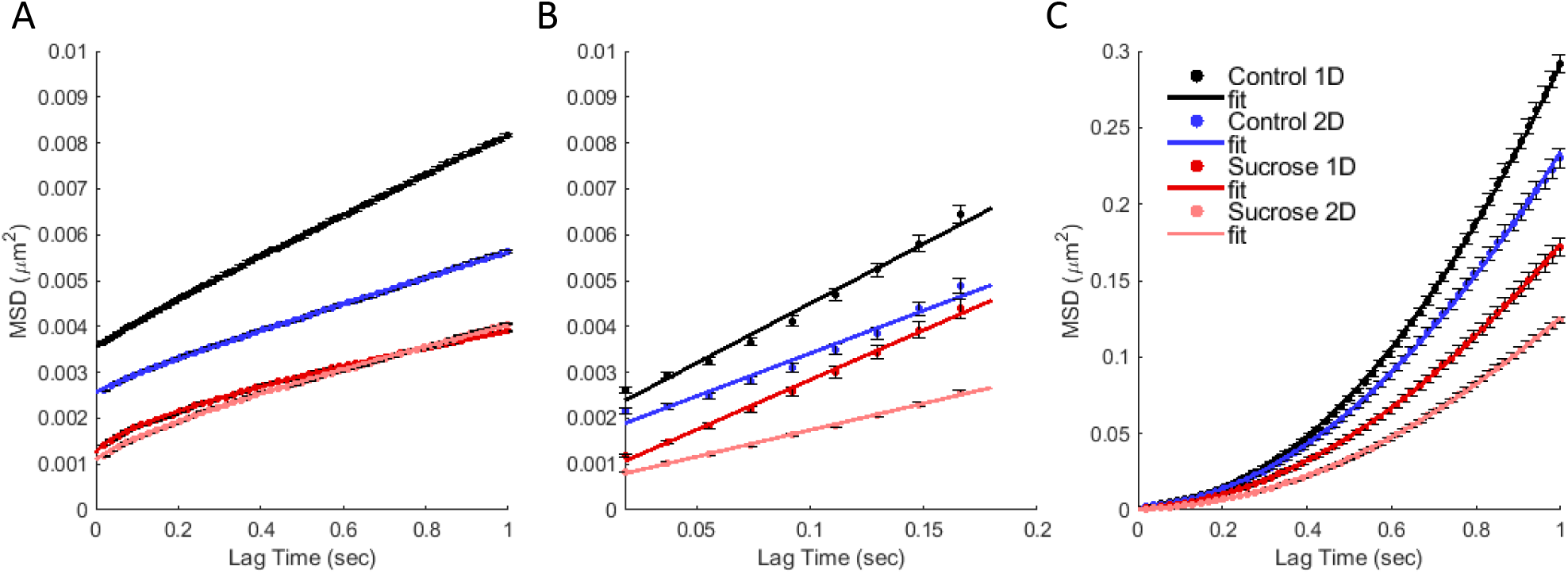
MSD per transport mode. (A) Sub-diffusive, (B) diffusive and (C) super-diffusive MSD plots of the respective lysosomal trajectories data points, as determined using the lMSD analysis. Color-coding indicates lysosomes inside aligned (1D) or randomly oriented (2D) neurites of differentiated PC12 cells in media without (control) or with sucrose.

Lysosomes exhibit a higher (vectorial) mean-squared displacement inside aligned neurites than in randomly oriented neurites, for all three motion modes studied. This alignment-associated effect appears to be consistent also in the case of the sucrose-induced perturbation, except for the sub-diffusive trajectory modes. As the resulting experimental values indicate (Table S.1), the diffusion coefficient of the diffusive trajectory modes is higher for lysosomes inside aligned neurites, without or with the presence of the perturbation (0.0065 *μm*^2^/*sec* and 0.0054 *μm*^2^/*sec*, respectively) as compared to lysosomes inside randomly oriented neurites (0.0047 *μm*^2^*/sec* and 0.0029 *μm*^2^/*sec*, respectively). Similarly, the drift velocity of the super-diffusive trajectory parts is higher for the lysosomes moving inside the aligned neurites, both in the media without and with sucrose (0.538 *μm/sec* and 0.397 *μm/sec*, respectively), in comparison to the respective values for randomly oriented neurites (0.460 *μm/sec* and 0.343 *μm/sec*, respectively). These findings are in agreement with the results for the x- and y-axis presented in Fig. 3, further confirming the association between neurite alignment and larger lysosomal displacements.

### Sucrose accumulation enhances crowding in aligned neurites

The experimental value of the sub-diffusive *α* exponent is found equal to 0.92 and 0.66 for lysosomes inside aligned neurites without and with sucrose, respectively. Thus, the decrease in the sub-diffusive *α* value, attributed to sucrose accumulation, is approximately equal to 0.26, that is twice the respective decrease observed inside randomly oriented neurites. The decrease in the diffusion coefficient, attributed to the incubation of the cells with sucrose, is approximately equal to 17% for lysosomes inside aligned neurites, almost half of the respective 35% for lysosomes inside randomly oriented neurites. Interestingly, our results indicate a sucrose-associated decrease in the drift velocity of the super-diffusive trajectory modes, approximately equal to 25%, which is the same for lysosomes inside both neurite configurations.

### Sucrose-induced lysosomal swelling results in proportionate decrease of the diffusion coefficient

Previously we found the mean diameter of lysosomes to be equal to < *d*_*c*_ >= 0.82 ± 0.02*μm* and < *d*_*s*_ >= 1.15 ± 0.02*μm* for differentiated PC12 cells in media without sucrose (control) versus in media with sucrose, respectively (Figure 2.E). According to the Stokes-Einstein equation (Albert 1905), the diffusion coefficient D of a spherical particle with radius *r*, through a liquid with low Reynolds number, is given by

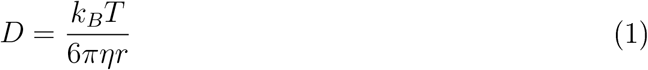

where *k*_*B*_ the Boltzmann constant, T the temperature and *η* the viscosity. Assuming the temperature and viscosity remain constant, we expect the ratio of two diffusion coefficients *D*_1_ and *D*_2_ of two particles to be analogous to the inverse ratio of their respective diameters *d*_1_ and 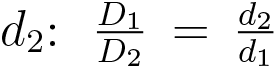. Thus, supposing there is no difference in the average temperature or viscosity of the cytoplasm between the cells in normal media and the cells in media containing sucrose, we expect the average diffusion coefficients and diameters of lysosomes in media without (control) versus with sucrose, to satisfy:

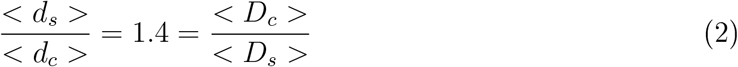

The experimental diffusion coefficients of the lysosomes inside cells in normal media and in sucrose-containing media, for the two configurations (Table S.1) yield: 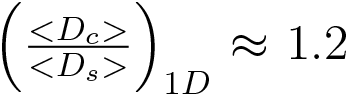 and 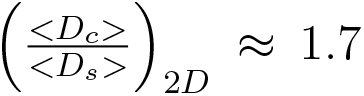, a result close to the expected value, with the case of the aligned configuration exhibiting lower deviation from the expected result.

### JDD analysis yields same characteristic values as MSD analysis for diffusive motion modes

The JDD is increasingly used for the characterization of intracellular trajectories (Ahmed and Saif 2014; Grünwald et al. 2008; Gal, Lechtman-Goldstein, and Weihs 2013). Thus, as a last step of our analysis, we investigated how the results of these distributions and the characteristic values extracted from their fits relate to the respective ones extracted from the MSD analysis.

In Fig. 6, the X- and Y- MSD curves of the diffusive and super-diffusive parts of lysosomal trajectories in neurites of differentiated PC12 cells for the four experimental conditions are displayed, along with the fitting curves. The respective probability distribution functions and corresponding fitting curves of the jump distances along the X- and Y- axis are presented in Fig. 7. Consistent with the MSD, the JD displacement along the X- axis (same as the neurite alignment axis) is larger in the case of the aligned neurites, for both without and with sucrose incubation conditions. Additionally, incubation with sucrose appears to be associated with smaller displacements, as compared to the control case.

**Figure 6:**
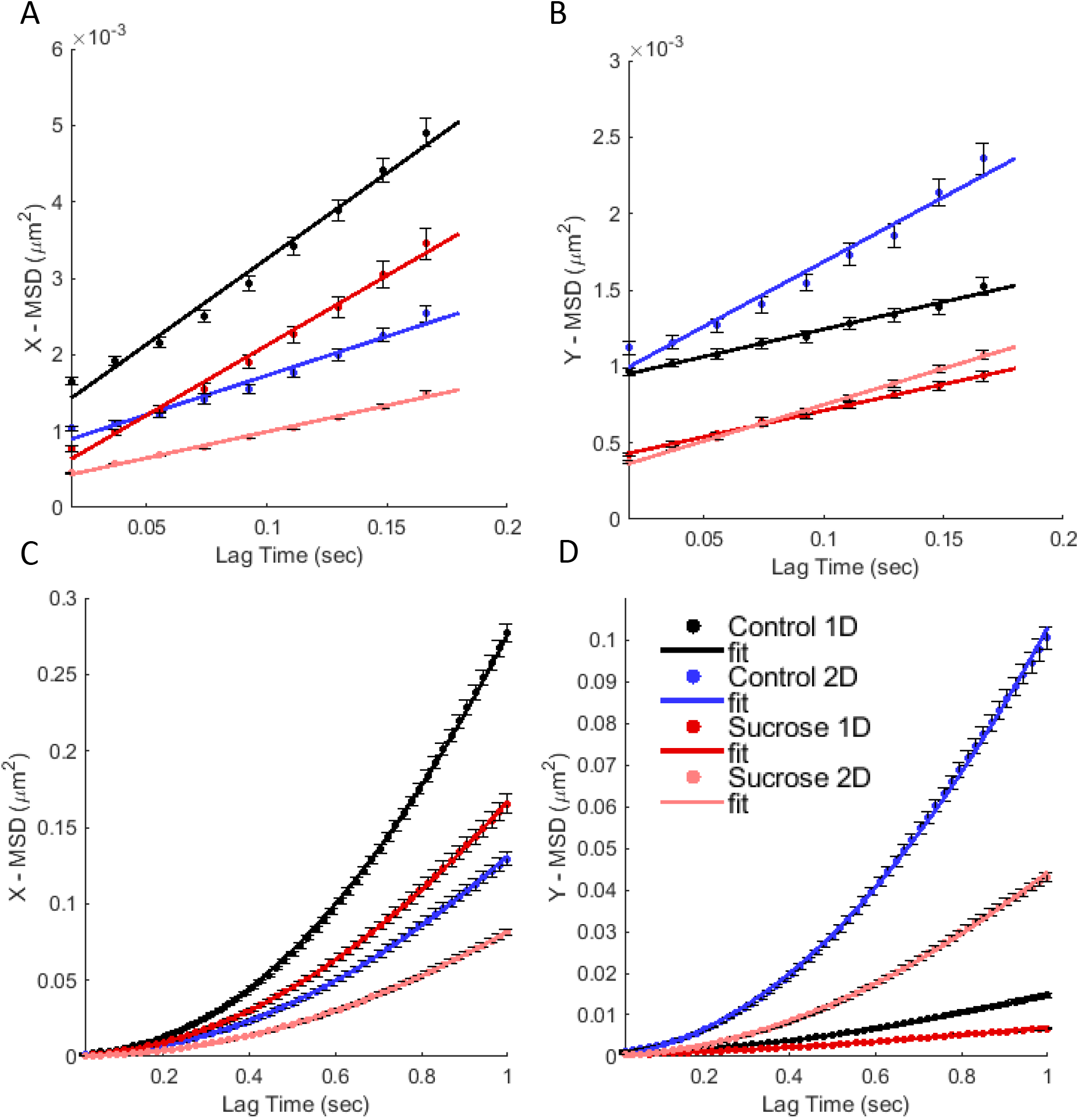
X- and Y- MSDs of diffusive and super-dffiusive trajectory parts. MSDs calculated collectively for all diffusive trajectory parts along (A) the X- and (B) Y- axis. MSDs calculated collectively for all super-diffusive trajectory parts along (C) the X- and (D) Y- axis. Color coding indicates data of lysosomes inside aligned (1D) and randomly oriented (2D) neurites of differentiatedPC12 cells in media without (control) or with sucrose. Diffusive MSDs (A, B) were fitted using equation 8. Super-diffusive MSDs (C, D) were fitted using equation 9. The fitting parameters are summarized in Table S.2.

**Figure 7:**
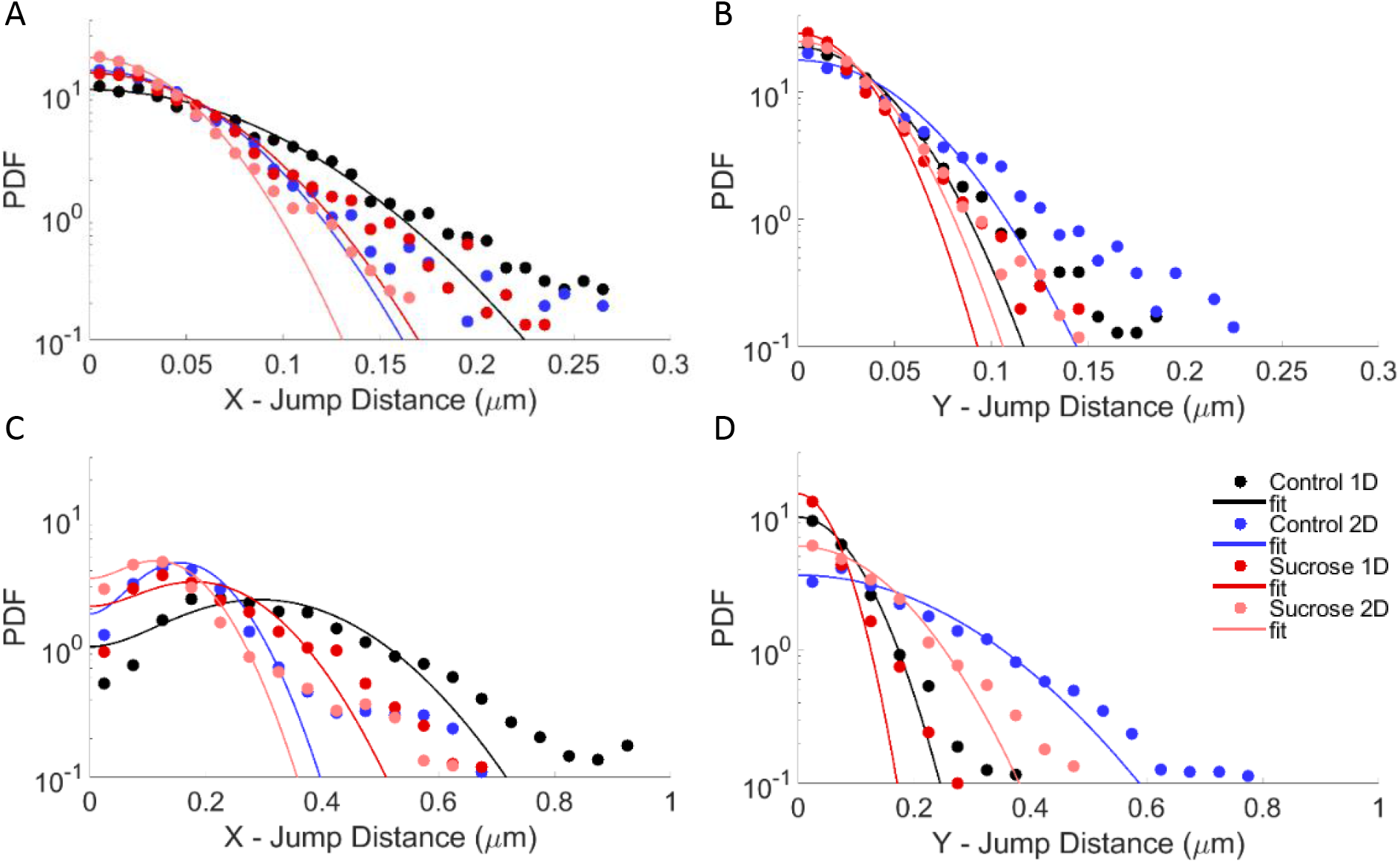
X- and Y- JDD PDFs of diffusive and super-diffusive trajectory parts. JDD PDFs calculated collectively for all diffusive trajectory parts along (A) the X- and (B) Y- axis, for *τ* = 240.5*ms*. JDD PDFs calculated collectively for all super-diffusive trajectory parts along (C) the X- and (D) Y- axis, for *τ* = 758.5*ms*. Color coding indicates data of lysosomes inside aligned (1D) and randomly oriented (2D) neurites of differentiated PC12 cells in media without (control) or with sucrose. Diffusive JDD PDFs (A, B) were fitted using equation 11. Super-diffusive JDD PDFs (C, D) were fitted using equation 12. The fitting parameters are summarized in Table S.3.

All diffusion coefficient and velocity values, resulting from the fits of the MSD and JDD curves are summarized in Table S.2 and S.3. It is remarkable how similar the resulting values of the X- and Y- component of the diffusion coefficient for the diffusive trajectory modes are. The resulting values of the X- component of the drift velocity of the super-diffusive trajectory modes exhibit small differences; the values extracted via fitting the JDD PDFs are systematically smaller than the respective ones derived from the MSD fits. This is of no surprise since, as can be seen, the fitting curves neglect the longer tails. However the trend observed among the experimental conditions is maintained: in the presence of sucrose the velocity is smaller than the control case, regardless the neurite configuration, and the 1D alignment indicates higher values than the random orientation case, regardless the presence of the perturbation. The Y- component of the drift velocity of the super-diffusive trajectory modes, estimated via fitting the JDD PDF, deviates from the other values by two orders of magnitude.

## Discussion

Cell survival, growth and conservation of homeostasis rest upon fine tuning and interplay of a plethora of processes. One such vital process is the transport of organelles, proteins or debris within the cellular environment. Malfunctioning intracellular motion is associated with neurodegenerative diseases (Aridor and Lisa A. Hannan 2000; Aridor and Lisa A Hannan 2002; Sleigh et al. 2019; Appert-Rolland, Ebbinghaus, and Santen 2013; De Vos and Hafezparast 2017), emphasizing the significance of this process for neuronal cells more than other cell types. However, a deeper insight is of essence, to determine whether faulty intracellular transport underlies or gives rise to the pathology of these diseases.

Here, we employ a model system of neuron-like cells (Lloyd A Greene and Tischler 1976) to characterize the motion of lysosomes inside their neurites. As the cell shape affects the organization of the cytoskeleton and thereby the intracellular transport, we investigate this effect by guiding the neurites of differentiated PC12 cells towards two distinct geometries: either randomly oriented on a surface, or aligned along chemically-patterned lines. In parallel, to mimic a pathological cellular phenotype, we perturb the cellular homeostasis via sucrose accumulation and induced lysosome swelling, and detect via motion analysis how its effect varies with the neurite geometry.

The overall MSD and JDD plots of lysosomal trajectories indicate enhanced transport when the neurites are aligned. The length scale of the observed trajectories is small, compared to the neurite width or curvature. Thus, this effect on the transport can be attributed to a global rearrangement of the cytoskeletal components, resulting from the alignment of the neurites, confirming the hypothesis that the cell shape impacts intracellular transport.

Implementing local MSD analysis, we separately characterize the sub-diffusive, diffusive and super-diffusive transport phases of the recorded lysosomal trajectories (Arcizet et al. 2008; Mahowald, Arcizet, and Heinrich 2009; Götz et al. 2015). We find that both diffusive and super-diffusive motion of lysosomes is enhanced inside aligned neurites. In addition, this result is maintained in the case where the homeostasis has been impaired via sucrose incubation, suggesting a global effect of neurite alignment on organelle transport. For the sub-diffusive trajectory parts, the difference is smaller, as seen from the alpha-exponent values of the MSD curves fits, which are slightly larger for lysosomes inside aligned neurites. Our findings complement previous studies, which used lMSD analysis to investigate the effect of cytoskeleton organization on intracellular dynamics of *Dictyostellium Discoideum* cells (Otten et al. 2012; Götz et al. 2015; Grady et al. 2017; Mahowald, Arcizet, and Heinrich 2009), demonstrating the potential of this analysis also for mammalian intracellular organelle motion.

In addition, we confirm that sucrose induces swelling of lysosomes inside differentiated PC12 cells with an associated increase of their average diameter by 0.33 *μ*m, and this leads to a proportionate decrease in their diffusion coefficient, as estimated by fitting the MSD curves of the diffusive trajectory modes. However, the neurite alignment seems to alleviate the sucrose effect in the case of the diffusive motion, resulting in half the respective decrease of the diffusion coefficient of lysosomes inside non-aligned neurites. Contrary to the findings reported by Bandyopadhyay et. al. (Bandyopadhyay et al. 2014), our analysis indicates a sucrose-associated decrease in the drift velocity of the super-diffusive transport modes, same for both neurite configurations. Furthermore, our results reveal a decrease in the alpha-exponent of the sub-diffusive trajectory modes, suggesting a crowding effect due to sucrose incubation (Weiss et al. 2004), more prominent inside aligned neurites.

Lastly, the JDD analysis of the diffusive motion modes resulted in values surprisingly close to the respective ones extracted from the MSD analysis. This was not the case for the super-diffusive trajectory parts, where the resulting values deviated significantly from the respective ones estimated using the MSD curve fitting. In future studies, it would be interesting to investigate whether this might be resolved after further discrimination of super-diffusive trajectory parts with *α* values between 1.1 and ~1.7 and active transport parts, with *α* values ~2 ±0.3, as the long tails in the JDD PDFs are neglected in the fits.

In summary, the experiments presented here are the first to quantitatively characterize the motion of functional organelles inside neurites of two different geometries, and compare the effect of sucrose-induced swelling on lysosomal motion, inside each neurite configuration and for each of the three transport modes. We confirm that the dimensionality of the neurites distinctly affects lysosome transport, with a positive correlation between 1D- neurite alignment, and higher diffusion coefficient and drift velocity of the lysosomal motion modes. Additionally we show that disruption of homeostasis via sucrose accumulation and induced lysosomal swelling, has larger effect when the motion occurs inside randomly oriented neurites. Our findings imply that, in physiological conditions, alterations in the ECM organization, which could for instance cause more branching of neurons’ axons, may enhance potential intracellular disrupted homeostasis, leading faster to pathological phenotypes. It would be interesting, in future research, to investigate the interplay between cell geometry and disease onset.

## Materials and Methods

### Laminin *μ*PIP

The mold for the patterning mask was fabricated with a Nanoscribe Photonic professional GT 3D laser printer (Nanoscribe, Germany), with two-photon polymerization (2PP) of IP-S photoresist (Nanoscribe n.d.). Prior to first use and after each subsequent use, a layer of trichloro(1H,1H,2H,2H-perfluorooctyl)silane, (Sigma-Aldrich) was deposited on the silicon mold (silanization), to reduce stiction (Srinivasan et al. 1998).

Poly dimethyl-siloxane (PDMS, Sylgard 184, Dow Corning, USA) was prepared by mixing the cross-linking agent with the elastomer base at a ratio of 1:10. The mixture was pipetted on the silicon mold and allowed to cross-link for 1 hour at 120*°*C. Subsequently, the hardened PDMS bearing the structure was peeled from the wafer.

For Plasma-Initiated Patterning, the PDMS mask was placed on an ibidi dish (ibidi GMBH, *μ*-Dish, 35 mm high, polymer coverslip bottom), with the structure-bearing side adherent to the bottom of the dish and was exposed to air plasma for 6 min 20 sec at 100 Watts (Diener Electronic Femto Plasma system).

Subsequently, the PDMS mask was removed and the substrate was flooded with 0.1% Pluronic F127 (Sigma-Aldrich) diluted in Phosphate Buffer Saline (PBS) for 45 minutes at room temperature. The dish was then washed 3 times with PBS and once with RPMI (Gibco™) and subsequently incubated with 25*μ*g/ml Laminin (Sigma-Aldrich) diluted in RPMI, for 1 hour at 37*°*C. Prior to seeding cells, the substrate was washed 3 times with RPMI.

### Cell Culture

PC12 cells (CH3 BioSystems) were cultured in dishes coated with rat-tail Collagen (CH3 BioSystems). Their growth medium consisted of 85% RPMI-1640 with Glutamax (Gibco−), 10% heat-inactivated Horse Serum (HS) (Sigma-Aldrich), 5% Fetal Calf Serum Heat Inactivated (FCS HI, Thermo Scientific) and 200*μ*g/mL pennicilin/streptomycin (PS). Media were refreshed three times per week, and the cells were split once per week at ratio 1:3-1:6. The cells were kept at 37*°*C and 5% CO_2_ in humidified atmosphere.

To induce differentiation, PC12 cells were seeded in ibidi uncoated dishes (ibidi GMBH, *μ*-Dish, 35 mm high) coated with Laminin (Sigma-Aldrich). For whole surface Laminin coating, the dish was exposed to air plasma for 6 min 20 sec at 100Watts (Diener Electronic Femto Plasma system). Subsequently, the dish was incubated for 1 hour at 37*°*C and 3% CO_2_ with 25 *μ*g/ml Laminin diluted in RPMI.

Cells were seeded at a density of 30.000 cells/cm^2^ in full media and after they had adhered, they were washed once with PBS and then the media were replaced with differentiation media consisting of Opti-MEM™ Reduced Serum Medium (Gibco™) supplemented with 0.5% Fetal Bovine Serum (Gibco™) and Nerve Growth Factor (NGF-2.5S Sigma-Aldrich) at final concentration of 100ng/ml. The differentiation media were refreshed three times per week.

To induce swelling of lysosomes, 50mM sucrose (Sigma-Aldrich) was added in the differentiation media 18 hours before imaging. At the end of the incubation, the cells were washed once with PBS and submerged again in normal culture or differentiation media.

Prior to imaging, the cells were incubated with 50-150nM Lysotracker (Invitrogen™) in RPMI for 30 minutes.

### Optical Microscopy

Optical microscopy images were acquired with a Nikon Ti Eclipse inverted microscope (NIKON corporation, Japan) equipped with a Yokogawa CSU-X1 spinning disc unit (10,000rpm, Andor Technology Ltd., United Kingdom). The samples were imaged with a 100x objective (Nikon CFI Plan Apo Lamda, NA 1.45). Excitation at 405nm, 488nm and 647nm was achieved via an Agilent MLC400 monolithic laser combiner (Agilent Technologies, Netherlands). The excitation light was filtered by a custom-made Semrock quad-band dichroic mirror for excitation wavelengths 400-410, 486-491, 460-570, and 633-647nm. The emitted light was filtered using a Semrock quad-band fluorescence filter (TR-F440-321-607-700), which has specific transmission bands at 440±40nm, 521±21nm, 607±34nm and 700±45nm and by Semrock Brightline single band fluorescence filters at 447±60nm (TR-F447-060) and 525±60nm (TR-F323-030). Images were captured with an Andor iXon Ultra 897 High-speed EM-CCD camera. Image acquisition was automated using NisElements software (LIM, Czech Republic). Time-lapse images were acquired every 18 ms, for up to 30 seconds. During data acquisition, the cells were kept in a humidified atmosphere at 37°C and supplied with 3% CO_2_ via the use of a Tokai Hit stage incubator.

### Data Analysis

The Feret diameter of lysosomes was calculated using FIJI (Schindelin et al. 2012). The selection of lysosomes in untreated cells was performed using an area mask between 0.1*μm*^2^ and 1.5*μm*^2^. Lysosomes in cells that had been treated with sucrose were selected with a mask of area between 0.5*μm*^2^ and 5.0*μm*^2^. Threshold values were the same for all images and clustered lysosomes were discarded from the analysis.

Trajectories of fluorescent lysosomes were tracked using the FIJI plugin, TrackMate (Tinevez et al. 2017), which returned the *x*− and *y*−coordinates of the center of the lysosomes, with a sub-pixel localization. During the tracking we included all three populations of lysosomes: those that appeared to be confined, diffusive and motile, with bigger (motor-mediated) displacements.

Further processing was performed using home-made Matlab algorithms. The *x*− and *y*−coordinates as a function of time for each trajectory were represented by a series of vectors at each time point *t*:

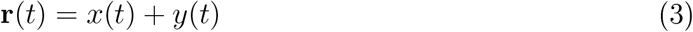

and the displacement Δ**r** at time *t* was calculatedas follows:

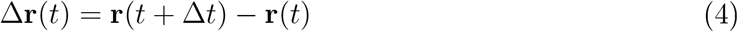

where Δ*t* is the inverse frame rate.

The Mean Squared Displacement (MSD) for lag time *τ* = *k*Δ*t* was calculated according to:

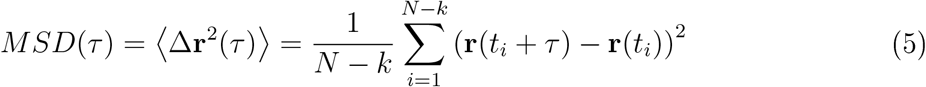

where *N* the number of data points in the trajectory and *k* = 1, 2, …, *N* − 1. The average MSD per condition is the average of the squared displacements of all lysosome trajectories for each lag time *τ*.

The local Mean Squared Displacement (lMSD) was calculated for each trajectory as described previously (Arcizet et al. 2008). Briefly, the MSD was calculated for each data point of the entire trajectory using a rolling window of 2.22 seconds (N=120, in eq. 5) and fitted for the interval 0-555ms (k=30 in eq. 5) with a power law:

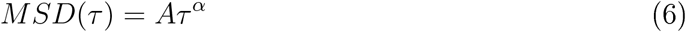

Tha alpha exponent as a function of time was subsequently used to partition the transport states as sub-diffusive for *α* < 0.9, diffusive for 0.9 ≤ *α* ≤ 1.1, or super-diffusive for *α* > 1.1.

In order to characterize more closely each type of motion, we analyzed collectively the respective trajectory parts, for each experimental condition (cells in media without (control) or with sucrose and in neurites that were aligned or randomly oriented).

The average MSD curve for each motion mode was calculated again according to eq. 5.

The sub-diffusive trajectory modes MSD was fitted with the power law describing anomalous diffusion

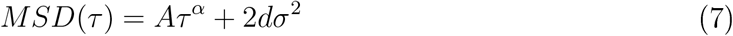

thereby obtaining the value of the anomalous *α* exponent (A is a constant). The MSD curve was fitted for all lag times (up to 30 sec).

The diffusive trajectory modes MSD was fitted using the equation describing Brownian motion

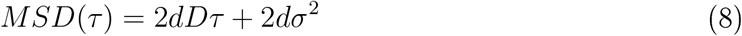

thus extracting the experimental value of the diffusion coefficient D. The MSD curve was fitted for lag times 1 to 10 (18.5-185 ms).

The super-diffusive trajectory modes MSD was fitted using the model of Brownian motion with drift (Briane, Vimond, and Kervrann 2019).

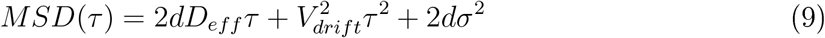

where *V*_*drift*_, the constant drift parameter models the velocity of the molecular motors. The MSD curve was fitted for lag times 1 to 55 (0.0185-1 sec).

In equations 7, 8 and 9 *σ* is the localization precision, *τ* the lag time and the parameter d refers to the dimensionality. d was set equal to 1 for the fit of the MSD along the x- or y- axis, and equal to 2 for the fit of the 2D- MSD curve.

The ump Distance Distribution (JDD) was calculated according to the self-part of the van Hove correlation function (Van Hove 1954):

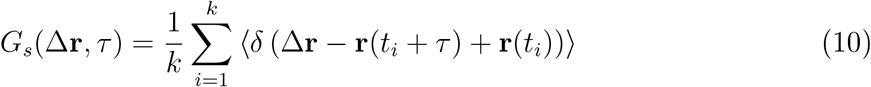

for displacements in both the *x*− and *y*− direction, and a specific lag time *τ*. *δ* here denotes the Dirac delta function in two dimensions and *k* = 1, 2, …, *N* − 1, with N the number of data points in the trajectory. The bin size was (arbitrarily) set to 1*μm* and the JDD was normalized into a probability density function.

The diffusive and super-diffusive trajectory parts for each experimental condition were used to calculate the respective JDD PDF, for lag times of 0.2405 ms and 0.7585 ms respectively. The PDFs were fitted to extract characteristic values of the motion, using the analytical expressions calculated in (Menssen and Mani 2019). Particularly, the diffusive trajectory modes JDD PDF was fitted for the x- direction, using

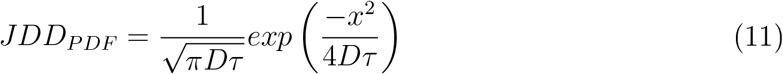

thereby estimating the experimental value of the diffusion coefficient D. Similarly, for the y-direction. The super-diffusive trajectory modes JDD PDF was fitted for the x- direction, using

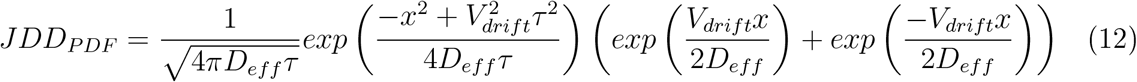

estimating the experimental value of the drift velocity *V*_*drift*_ and effective diffusion coefficient *D*_*eff*_. Likewise for the y-direction.

## Supplenentary Infornation

**Table S.1:** Motion parameters resulting from fitting the sub-diffusive, diffusive and super-diffusive trajectory parts MSD curves (demonstrated in Fig. 5), using eq. 7, 8, and 9, respectively.

| Motion mode | Sub-diffusive | Diffusive | Super-diffusive |  |
| --- | --- | --- | --- | --- |
| | $\alpha$ exponent | D ($\mu m^2/sec$) | $V_{drift}$ ($\mu m/sec$) | $D_{eff}$ ($\mu m^2/sec$) |
| Control 1D | $0.92 \pm 0.01$ | $0.0065 \pm 0.0013$ | $0.538 \pm 0.008$ | $5E - 8 \pm 0.002$ |
| Control 2D | $0.90 \pm 0.03$ | $0.0047 \pm 0.0015$ | $0.460 \pm 0.010$ | $0.005 \pm 0.002$ |
| Sucrose 1D | $0.66 \pm 0.03$ | $0.0054 \pm 0.0007$ | $0.397 \pm 0.004$ | $0.004 \pm 0.001$ |
| Sucrose 2D | $0.77 \pm 0.03$ | $0.0029 \pm 0.0003$ | $0.343 \pm 0.006$ | $0.002 \pm 0.001$ |
| | $\sigma$ ($\mu m$) | $\sigma$ ($\mu m$) | $\sigma$ ($\mu m$) | |
| Control 1D | $0.030 \pm 0.003$ | $0.022 \pm 0.012$ | $0.022 \pm 0.022$ | |
| Control 2D | $0.025 \pm 0.003$ | $0.020 \pm 0.012$ | $0.016 \pm 0.028$ | |
| Sucrose 1D | $0.018 \pm 0.004$ | $0.013 \pm 0.009$ | $0.015 \pm 0.014$ | |
| Sucrose 2D | $0.016 \pm 0.004$ | $0.012 \pm 0.005$ | $0.011 \pm 0.015$ | |

**Table S.2:**
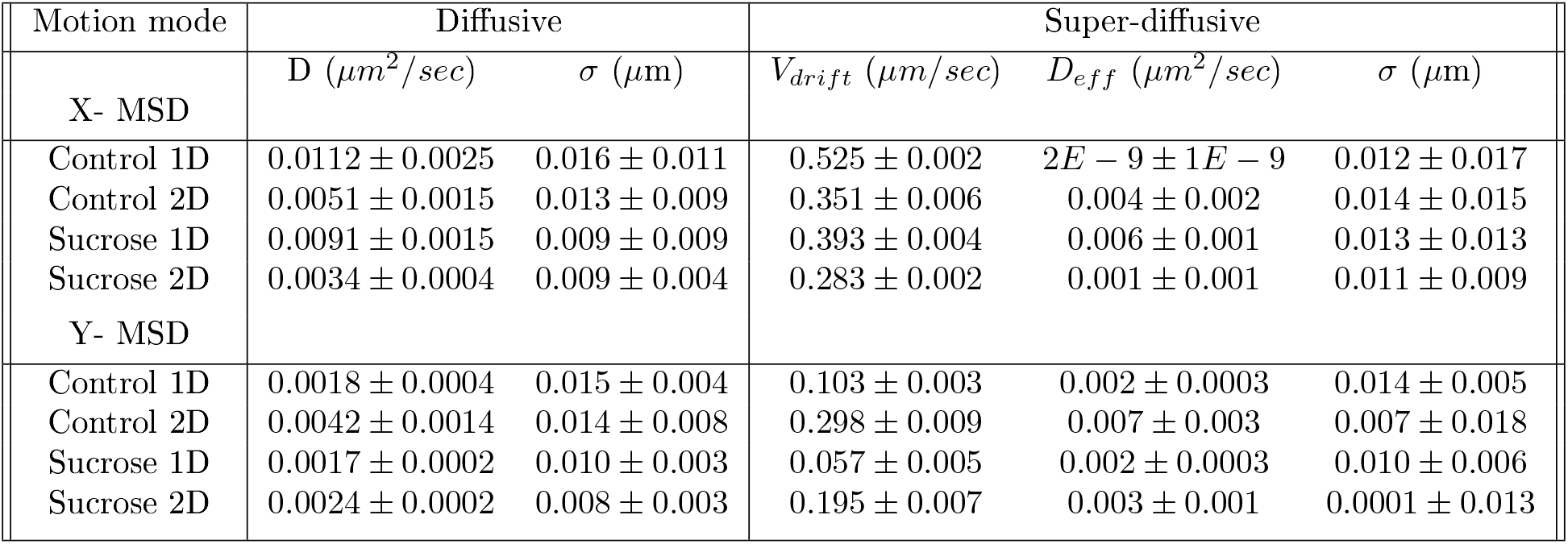
Motion parameters resulting from fitting the diffusive and super-diffusive trajectory parts x- and y- MSD curves (demonstrated in Fig. 6), using eq. 8 and 9.

| Motion mode | Diffusive |  | Super-diffusive |  |  |
| --- | --- | --- | --- | --- | --- |
| | D ($\mu m^2/sec$) | $\sigma$ ($\mu m$) | $V_{drift}$ ($\mu m/sec$) | $D_{eff}$ ($\mu m^2/sec$) | $\sigma$ ($\mu m$) |
| X- MSD |  |  |  |  |  |
| Control 1D | $0.0112 \pm 0.0025$ | $0.016 \pm 0.011$ | $0.525 \pm 0.002$ | $2E - 9 \pm 1E - 9$ | $0.012 \pm 0.017$ |
| Control 2D | $0.0051 \pm 0.0015$ | $0.013 \pm 0.009$ | $0.351 \pm 0.006$ | $0.004 \pm 0.002$ | $0.014 \pm 0.015$ |
| Sucrose 1D | $0.0091 \pm 0.0015$ | $0.009 \pm 0.009$ | $0.393 \pm 0.004$ | $0.006 \pm 0.001$ | $0.013 \pm 0.013$ |
| Sucrose 2D | $0.0034 \pm 0.0004$ | $0.009 \pm 0.004$ | $0.283 \pm 0.002$ | $0.001 \pm 0.001$ | $0.011 \pm 0.009$ |
| Y- MSD |  |  |  |  |  |
| Control 1D | $0.0018 \pm 0.0004$ | $0.015 \pm 0.004$ | $0.103 \pm 0.003$ | $0.002 \pm 0.0003$ | $0.014 \pm 0.005$ |
| Control 2D | $0.0042 \pm 0.0014$ | $0.014 \pm 0.008$ | $0.298 \pm 0.009$ | $0.007 \pm 0.003$ | $0.007 \pm 0.018$ |
| Sucrose 1D | $0.0017 \pm 0.0002$ | $0.010 \pm 0.003$ | $0.057 \pm 0.005$ | $0.002 \pm 0.0003$ | $0.010 \pm 0.006$ |
| Sucrose 2D | $0.0024 \pm 0.0002$ | $0.008 \pm 0.003$ | $0.195 \pm 0.007$ | $0.003 \pm 0.001$ | $0.0001 \pm 0.013$ |

**Table S.3:** Motion parameters resulting from fitting the diffusive and super-diffusive trajectory parts x- and y- JDD PDFs (demonstrated in Fig. 7), using eq. 11 and 12.

| Motion mode | Diffusive | Super-diffusive |  |
| --- | --- | --- | --- |
| | D ($\mu m^2/sec$) | $V_{drift}$ ($\mu m/sec$) | $D_{eff}$ ($\mu m^2/sec$) |
| X - JDD |  |  |  |
| Control 1D | $0.0112 \pm 0.0016$ | $0.388 \pm 0.060$ | $0.019 \pm 0.009$ |
| Control 2D | $0.0054 \pm 0.0006$ | $0.205 \pm 0.024$ | $0.005 \pm 0.002$ |
| Sucrose 1D | $0.0060 \pm 0.0007$ | $0.243 \pm 0.055$ | $0.010 \pm 0.007$ |
| Sucrose 2D | $0.0033 \pm 0.0003$ | $0.158 \pm 0.026$ | $0.005 \pm 0.002$ |
| Y - JDD |  |  |  |
| Control 1D | $0.0026 \pm 0.0003$ | $0.001 \pm 17931$ | $0.004 \pm 12.785$ |
| Control 2D | $0.0042 \pm 0.0009$ | $0.007 \pm 5425$ | $0.032 \pm 30.580$ |
| Sucrose 1D | $0.0016 \pm 0.0007$ | $0.001 \pm 3958$ | $0.002 \pm 3.556$ |
| Sucrose 2D | $0.0021 \pm 0.0002$ | $0.001 \pm 79828$ | $0.012 \pm 78.405$ |

**Table S.4:**
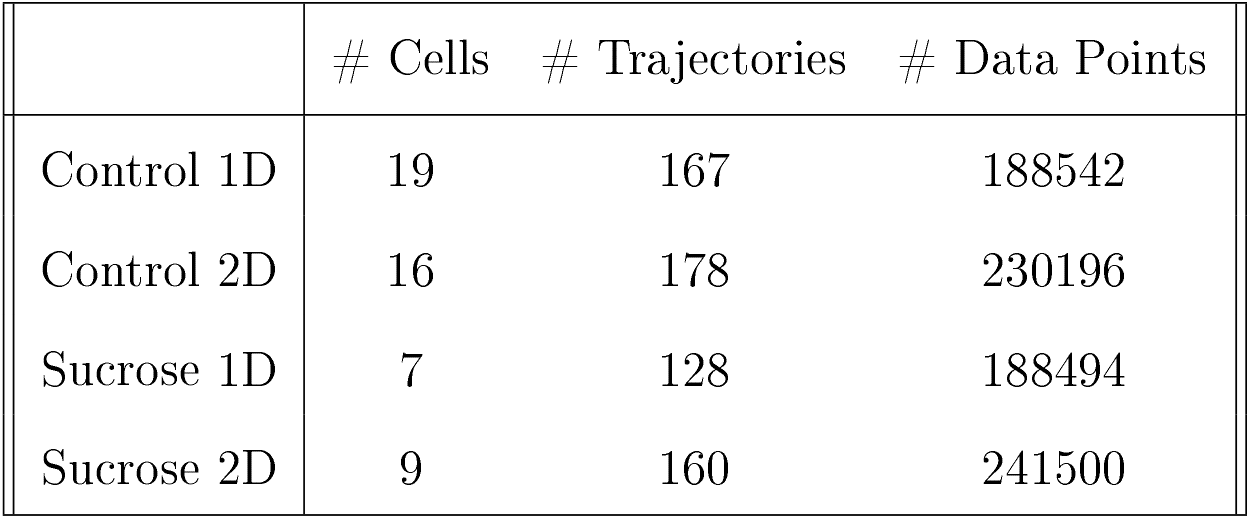
Data Statistics. Number of cells imaged, number of total lysosomal trajectories tracked and number of total data points per experimental condition (differentiated PC12 cells in media without (control) and with sucrose, with their neurites aligned (1D) or randomly oriented (2D)).

|  | # Cells | # Trajectories | # Data Points |
| --- | --- | --- | --- |
| Control 1D | 19 | 167 | 188542 |
| Control 2D | 16 | 178 | 230196 |
| Sucrose 1D | 7 | 128 | 188494 |
| Sucrose 2D | 9 | 160 | 241500 |

**Table S.5:** Data Statistics of trajectories modes. Number of trajectory modes and number of total data points falling for each transport category (sub-diffusive, diffusive and super-diffusive), as determined with lMSD analysis.

|  | Sub-diffusive |  | Diffusive |  | Super-diffusive |  |
| --- | --- | --- | --- | --- | --- | --- |
|  | # Modes | # Data Points | # Modes | # Data Points | # Modes | # Data Points |
| Control 1D | 352 | 154732 | 334 | 5484 | 135 | 8254 |
| Control 2D | 307 | 196326 | 257 | 4504 | 117 | 8029 |
| Sucrose 1D | 365 | 154513 | 426 | 7110 | 185 | 10941 |
| Sucrose 2D | 600 | 187570 | 777 | 14336 | 323 | 20111 |

## Acknowledgnents

The authors would like to acknowledge the Fraunhofer Gesellschaft for the Fraunhofer Attract grant “3D NanoCell”.

